# GNMFLMI: Graph Regularized Nonnegative Matrix Factorization for Predicting LncRNA-MiRNA Interactions

**DOI:** 10.1101/835934

**Authors:** Mei-Neng Wang, Zhu-Hong You, Li-Ping Li, Leon Wong, Zhan-Heng Chen, Cheng-Zhi Gan

**Affiliations:** School of Mathematics and Computer Science, Yichun University, Yichun Jiangxi 336000, China; Xinjiang Technical Institutes of Physics and Chemistry, Chinese Academy of Sciences, Urumqi 830011, China

**Keywords:** lncRNA-miRNA interaction, nonnegative matrix factorization, graph regularization, lncRNA-miRNA similarity

## Abstract

Long non-coding RNAs (lncRNAs) and microRNAs (miRNAs) have been involved in various biological processes. Emerging evidence suggests that the interactions between lncRNAs and miRNAs play an important role in regulating of genes and the development of many diseases. Due to the limited scale of known lncRNA-miRNA interactions, and expensive time and labor costs for identifying them by biological experiments, more accurate and efficient lncRNA-miRNA interactions computational prediction approach urgently need to be developed. In this work, we proposed a novel computational method, GNMFLMI, to predict lncRNA-miRNA interactions using graph regularized nonnegative matrix factorization. More specifically, the similarities both lncRNA and miRNA are calculated based on known interaction information and their sequence information. Then, the affinity graphs for lncRNAs and miRNAs are constructed using the *p*-nearest neighbors, respectively. Finally, a graph regularized nonnegative matrix factorization model is developed to accurately identify potential interactions between lncRNAs and miRNAs. To evaluate the performance of GNMFLMI, five-fold cross validation experiments are carried out. GNMFLMI achieves the AUC value of 0.9769 which outperforms the compared methods NMF and CNMF. In the case studies for lncRNA nonhsat159254.1 and miRNA hsa-mir-544a, 20 and 16 of the top-20 associations predicted by GNMFLMI are confirmed, respectively. Rigorous experimental results demonstrate that GNMFLMI can effectively predict novel lncRNA-miRNA interactions, which can provide guidance for relevant biomedical research.

## 1 Introduction

With the development of next-generation sequencing, specific biological mechanisms can be better understood from the wide-ranging biomolecular interactions in the genome. Long non-coding RNAs (lncRNAs) and microRNAs (miRNAS) were previously thought to be non-functional sequences in the process of gene evolution [1]. In fact, they not only play an important role in cell differentiation, somatic development and other life processes, but also can participate in the occurrence of disease through interaction [1]. LncRNA is a type of non-coding RNA (ncRNA) located in the nucleus or cytoplasm of more than 200nt in length which has no obvious protein-coding function and exists in any branch of life [2]. Depending on the positional relationship of the coding genes, lncRNAs can be divided into five categories (i.e. bidirectional, antisense, sense, introverted and intergenic) [3–5]. Because of lncRNA has tissue specificity, cell specificity, spatiotemporal specificity, developmental stage specificity and disease specificity, it is widely involved in cell differentiation, metabolism and proliferation, and is closely associated with many complex diseases [6, 7]. More and more evidences have shown that lncRNAs can silence or activate genes by regulating histone modification, DNA methylation, mRNA splicing and chromatin remodeling in a variety of ways, such as epigenetics, transcriptional regulation, and post-transcriptional control and so on [8]. As a new focus of regulation for gene expression, lncRNA plays a biological role mainly through signal function, bait function, scaffold function and guiding function [9]. Even though the experiment has identified more than 58 000 human lncRNA genes, Only a few lncRNAs have been functionally characterized, such as H19, HOTAIR and Malat, Most of them are still functionally uncharacterized [10].

LncRNAs participate in the regulation of expressed proteins via specific mechanism involving multiple biological interactions such as lncRNA-mRNA, lncRNA-ncRNA and lncRNA-protein interactions [11]. Therefore, it is necessary to construct a network of biomolecular interactions mediated by lncRNAs, which is very useful for revealing the underlying mechanisms and biological functions of lncRNAs [12]. As a bait for miRNA, lncRNAs can inhibit the binding of miRNA to target gene mRNA, and can also act as an endogenous miRNA sponge to inhibit miRNA expression [13, 14]. With the accumulation of knowledge on miRNA function, the lncRNA-miRNA interaction network can help us better understand the complex functions of lncRNA [7]. MiRNA is a class of non-coding short sequence RNAs of 18-25nt in length that are widely found in eukaryotes and are highly conservative, spatiotemporal-specific and tissue-specific [15]. A miRNA molecule can regulate the expression of up to 200 target genes, and about one-third of human genes are regulated by miRNAs [15]. Up to now, miRNAs are considered to be the most important gene regulators in cell differentiation, development, growth, and tumorigenesis, progression, metastasis, and drug resistance [16]. In the occurrence and development of human tumors, some miRNAs can act as both an oncogenes and a cancer suppressor genes [17, 18]. For example, certain miRNAs are associated with the development of ductal carcinoma in situ (DCIS) to invasive carcinoma, especially miR-210, miR221 and let-7d, which are down-regulated in situ carcinoma but up-regulated in invasive carcinoma [17].

In recent years, More and more studies have shown that both lncRNA and miRNA play critical roles in various biological processes and human complex diseases [19, 20]. It has been systematically studied that the lncRNA-miRNA interactions exert regulation role in some human complex diseases [21–23]. In many diseased cells, lncRNA is discovered to have a certain quantitative relationship with some miRNAs, this quantitative relationship is closely associated with the occurrence and development of diseases [24]. For example, in the renal cell carcinoma (RCC), after Malta silencing, miR-205 expression is up-regulated, but cell proliferation, migration and invasion are inhibited. Conversely, the expression of Malta is significantly reduced after overexpression of miR-205, experiments showed that there is a mutual regulation relationship between Malta and miR-205 [25]. The detailed understanding of interactions between lncRNAs and miRNAs in disease is very helpful for new biomarker discovery and treatment methods exploration [26]. However, identifying the interactions between lncRNAs and miRNAs is expensive and time-consuming by biological experiment.

To accelerate the process of identifying interactions between biomolecules, many computational methods have been proposed and effectively used for predicting relationships (e.g. miRNA-disease associations, protein-protein interactions and lncRNA-protein interactions), including manifold learning, manifold embedding and semi-supervised learning, etc. [27–29]. The computational methods for predicting miRNA-target interactions usually have the following common rules, including site accessibility, seed matching, free energy and protection [10, 30]. However, many miRNA-target identification methods were proposed originally for mRNA targets that may not be able to identify the interactions between lncRNAs and miRNAs, or even contradictory [31, 32]. Huang and Chan proposed the EPLMI calculation model based on the assumption that lncRNAs tend to interact collaboratively with miRNAs of similar expression profiles, and constructs bipartite graph via known lncRNA-miRNA interactions for prediction[33]. Huang *et al.* proposed the GBCF computational model based on known interaction network to obtain a top*-k* probability ordering list of individual lncRNA or miRNA for prediction [34]. Although the above two methods have better predictive effects in the known lncRNA-miRNA interaction network, they cannot be applied to new lncRNA or miRNA. The predictive effect of interactions between molecules can be improved by integrating biological information from different sources [35–37]. In fact, prediction of lncRNA-miRNA interactions can be considered as a recommender system problem [38, 39]. Accumulated studies have shown that matrix factorization is an effective method which has been successfully used in recommender system for data representation, and already widely applied in the field of bioinformatics [40–42].

In this paper, we propose a new calculation model, GNMFLMI, to predict lncRNA-miRNA interactions using graph regularized nonnegative matrix factorization (NMF) [43]. The model is based on the assumption that functionally similar lncRNAs (or miRNAs) are more possibility to interact with a same miRNA (or lncRNA) [44]. GNMFLMI fully exploits miRNA/lncRNA sequence information and known lncRNA-miRNA interaction network to calculate miRNA/lncRNA similarity. Subsequently, the graph spaces of miRNA and lncRNA were constructed based on the local invariance hypothesis of intrinsic geometric space [45–47], which promote similar lncRNAs/miRNAs to be close enough to each other in the lncRNA/miRNA space. We evaluated the performance of our method by five-fold cross validation and compared the performance with standard NMF [43], RNMF [48]. The experiment results show that GNMFLMI is better than other compared methods, and can effectively predict the novel lncRNA-miRNA interactions.

## 2 Materials and methods

### 2.1 Benchmark Dataset

The lncRNA-miRNA interactions dataset used in this work were obtained from the lncRNASNP2 database in January, 2019. This dataset is collected and collated by Ya-Ru Miao *et al.* [49] and is accessible to academic users at http://bioinfo.life.hust.edu.cn/lncRNASNP. We downloaded the known lncRNA-miRNA interactions and the duplicated entries were removed. After the preprocessing, 8634 experimentally verified lncRNA–miRNA interactions were obtained, containing 262 miRNAs and 468 lncRNAs. In the experiment, all known lncRNA-miRNA interactions provided by lncRNASNP2 dataset were used as positive samples, and other unknown lncRNA-miRNA interactions as negative samples. The lncRNA-miRNA interactions adjacency matrix *Y* ∈ *R*^*r*×*n*^ was constructed based on lncRNASNP2 database, where *r* is the number of lncRNAs, *n* is the number of miRNAs. If the lncRNA *l*(*i*) was verified to be interacted with miRNA *m*(*j*), the element *Y*(*i*, *j*) was assigned the value of 1, otherwise it is 0.

In this study, we let *L* = {*l*_1_, *l*_2_, ⋯, *l*_*r*_} and *M* = {*m*_1_, *m*_2_, ⋯, *m*_*n*_} which denote the set of *r* lncRNAs and *n* miRNAs. The *ith* row vector of matrix *Y*, *Y*(*l*_*i*_) = (*Y*_*i*1_, *Y*_*i*2_, ⋯, *Y*_*in*_); the *jth* column vector of matrix *Y*, *Y*(*m*_*j*_) = (*Y*_1*j*_, *Y*_2*j*_, ⋯, *Y*_*rj*_). *Y*(*l*_*i*_) and *Y*(*m*_*j*_) represent the interaction profiles for lncRNA *l*_*i*_ and miRNA *m*_*j*_, respectively.

### 2.2 Related work

#### 2.2.1 The standard nonnegative matrix factorization (NMF)

Nonnegative matrix factorization (NMF) is an effective algorithm which has been successfully used in recommender system for data representation [41]. This algorithm divides a large original matrix *Y* into two low-dimensional nonnegative matrices: a basis matrix *U* and a coefficient matrix *V*, and make their product as close as possible to the original matrix. Recent years, NMF has been successfully utilized to predict potential associations of lncRNA-protein [50], miRNA-disease [51], Microbe-disease [52], drug-target [53], CircRNA-disease [54], etc. In this work, the lncRNA-miRNA interaction adjacency matrix *Y* ∈ *R*^468×262^ is divided into *U* ∈ *R*^*k*×468^ and *V* ∈ *R*^*k*×262^, *k* is the sub-space dimensionality (*k* < *rn*/(*r* + *n*)). So that:

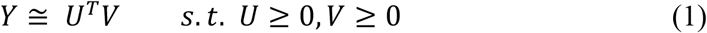

The objective function for predicting lncRNA-miRNA interactions can be formulated as following:

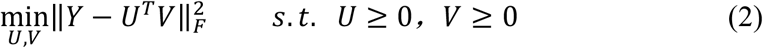

Where, ||∙||_*F*_ denotes the Frobenius norm. *U*, *V* ≥ 0 means that all elements of *U* and *V* are nonnegative. According to matrix properties 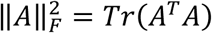, 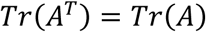, and 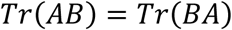, we can obtain:

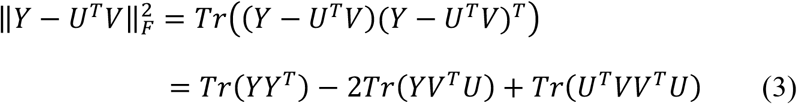

Where *Tr*(∙) is the trace of a matrix.

Lee and Seung [43] propose the nonnegative matrix factorization algorithm and this algorithm is based on the multiplicative update rules of *U* and *V*, the update rules are as follows:

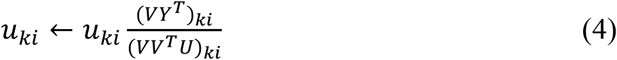

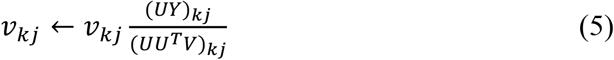

#### 2.2.2 Constrained nonnegative matrix factorization (CNMF)

Due to the standard NMF cannot ensure the *U* and *V* smoothness. We can use the Tikhonov (*L*_2_) in the standard NMF to solve this problem [55]. Pauca *et al.* proposed the following constrained nonnegative matrix factorization (CNMF) formulation [48]:

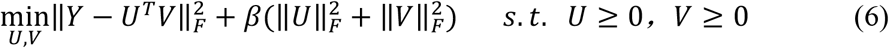

where *β* is the sparseness constraint coefficient which is used adjust the sparsity of *U* and *V* via *L*_2_-norm. The Eq. (6) can be written as:

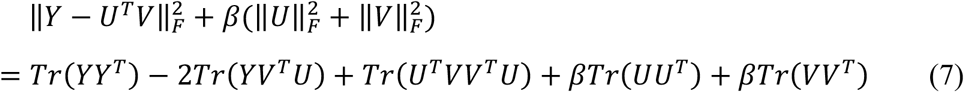

The multiplicative update rules of *U* and *V* are as follows:

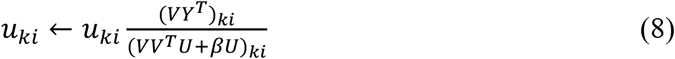

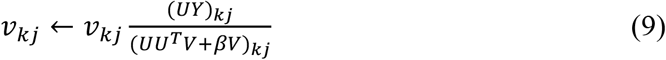

the details of multiplicative update rules are in section 2.3.5.

### 2.3 Graph regularized nonnegative matrix factorization for predicting lncRNA-miRNA interactions (GNMFLMI)

#### 2.3.1 methods overview

In this study, we propose a new calculation model, GNMFLMI, to predict lncRNA-miRNA interactions. The GNMFLMI can be summarized into three steps, and its framework is shown in Figure 1. First, the similarity matrices of lncRNA and miRNA are calculated based on lncRNA/miRNA sequence information and known lncRNA-miRNA interaction network. Second, we construct the affinity graphs for lncRNAs and miRNAs using *p*-nearest neighbors. Finally, graph regularized nonnegative matrix factorization is performed to calculate the lncRNA-miRNA interaction scores.

**Fig. 1.**
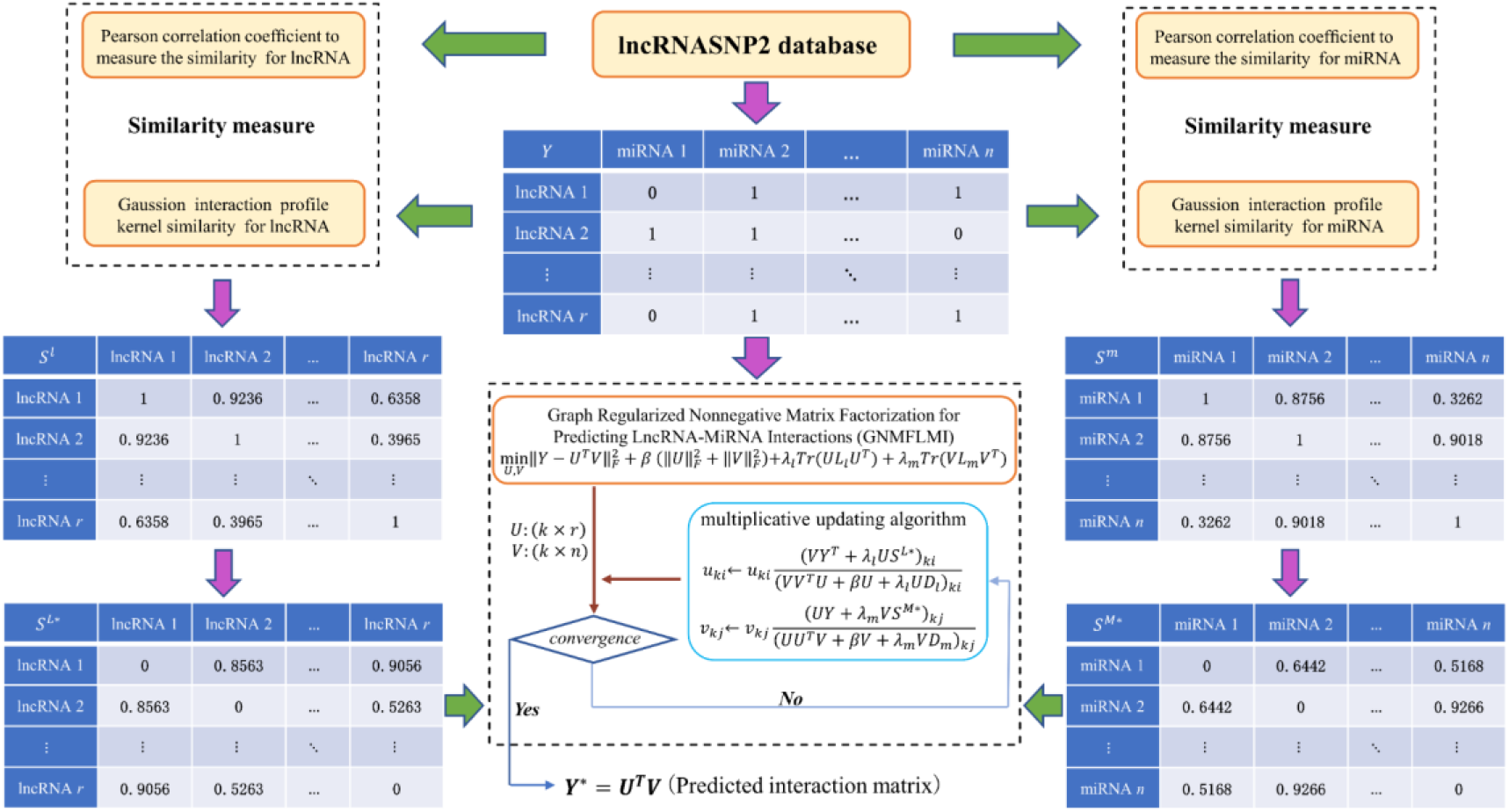
Flowchart of GNMFLMI for predicting potential lncRNA-miRNA interaction.

#### 2.3.2 Similarity measure

In our method, we need to construct the similarity matrices for lncRNAs and miRNAs separately, so the similarity both each pair of lncRNA-lncRNA and each pair of miRNA-miRNA need to be determined. In this study, two different types of lncRNA/miRNA similarity were measured via integrating diverse sources of information. The first type of lncRNA/miRNA similarity is calculated using Gaussian interaction profile (GIP) kernel based on known lncRNA-miRNA interaction network [56]. The second type is calculated using Pearson correlation coefficient (PCC) based on lncRNA/miRNA sequence information [57]. Subsequently, the overall similarity matrices for lncRNAs and miRNAs were constructed based on above two types of similarity.

##### Similarity based on Gaussian interaction profile (GIP) kernel

According to the assumption that functionally similar lncRNAs tend to interact with the similar miRNAs, and Gaussian interaction profile (GIP) kernel has been widely used to compute the molecule similarity [56, 58]. The Gaussian interaction profile kernel similarity of lncRNA and miRNA can be constructed according to the topologic information of known lncRNA-miRNA interaction network. Thus, we use GIP kernel to calculate the similarity *L*_*GIP*_(*l*_*i*_, *l*_*j*_) between lncRNA *l*_*i*_ and lncRNA *l*_*j*_ as following:

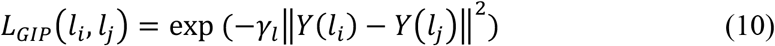

where

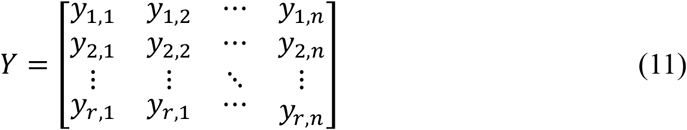

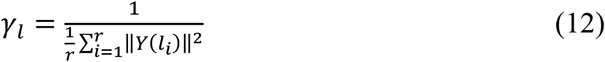

*Y* is the adjacent matrix of lncRNA-miRNA interaction based on lncRNASNP2 database. *r* and *m* are the number of lncRNAs and miRNAs, respectively. The size of *L*_*GIP*_ is *r* × *r*, *Y*(*l*_*i*_) is the *ith* row of the adjacent matrix *Y*, *γ*_*l*_ is the kernel width parameter.

Similar to lncRNAs, the Gaussian interaction profile kernel similarity *M*_*GIP*_(*m*_*i*_, *m*_*j*_) of miRNA *m*_*i*_ and miRNA *m*_*j*_ can be calculated as follows:

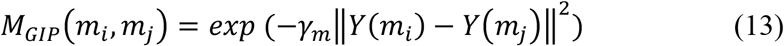

where

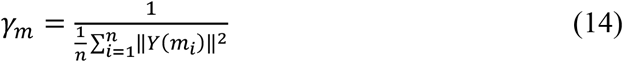

The size of *M*_*GIP*_ is *n* × *n*, *Y*(*m*_*i*_) is the *ith* column of the adjacent matrix *Y*, *γ*_*m*_ is the kernel width parameter.

##### Similarity based on Pearson correlation coefficient (PCC)

We download the expression profiles for lncRNAs and miRNAs from lncRNASNP2 database. For each lncRNA/miRNA, the values of expression profile can be obtained. Pearson correlation coefficient (PCC) has been widely applied to study expression profiles in bioinformatics [59, 60]. PCC of lncRNA/miRNA is calculated based on the lncRNA/miRNA expression profile values. For example, Given two lncRNAs *l*_*i*_ and *l*_*j*_, the expression profiles are denoted as *X*_*l*_ = {*x*_*l*1_, *x*_*l*2_, ⋯, *x*_*lt*_} and *Z*_*l*_ = {*z*_*l*1_, *z*_*l*2_, ⋯, *z*_*lt*_}. The similarity between lncRNA *l*_*i*_ and lncRNA *l*_*j*_ is calculated as follows:

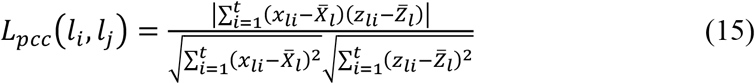

where, *t* is the number of attributes of the expression profile, 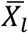 and 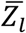 denote the average value of *X*_*l*_ and *Z*_*l*_, respectively. Generally, the larger *L*_*pcc*_(*l*_*i*_, *l*_*j*_) represents the more similarly expression of lncRNA *l*_*i*_ and lncRNA *l*_*j*_.

Similar to lncRNAs, the similarity between miRNA *m*_*i*_ and miRNA *m*_*j*_ can be calculated by Pearson correlation coefficient as follows.

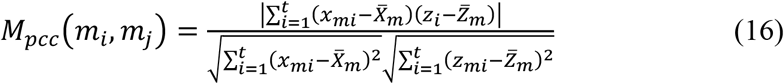

where, *X*_*m*_ = {*x*_*m*1_, *x*_*m*2_, ⋯, *x*_*mt*_} and *Z*_*m*_ = {*z*_*m*1_, *z*_*m*2_, ⋯, *z*_*mt*_} denote the expression profiles of miRNA *m*_*i*_ and miRNA *m*_*j*_. 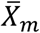 and 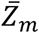 denote the average value of *X*_*m*_ and *Z*_*m*_, respectively.

##### Construct the overall similarity for lncRNAs and miRNAs

In this study, the Gaussian interaction profile kernel similarity and Pearson correlation coefficient similarity of lncRNA and miRNA are calculated, respectively. After that, the functional similarity between lncRNA *l*_*i*_ and lncRNA *l*_*j*_ is defined according to [61, 62], the final similarity matrix *S*^*L*^ of lncRNA is calculated as follows:

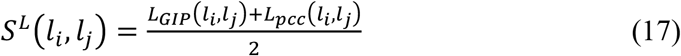

where *S*^*L*^ is *r*-order square matrix, *S*^*L*^(*l*_*i*_, *l*_*j*_) represents the similarity score between lncRNA *l*_*i*_ and lncRNA *l*_*j*_.

Based on the same method, the final similarity matrix *S*^*M*^ of miRNA is calculated as follows:

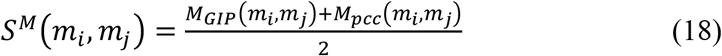

where *S*^*M*^ is *n*-order square matrix, *S*^*M*^(*m*_*i*_, *m*_*j*_) represents the similarity score between miRNA *m*_*i*_ and miRNA *m*_*j*_.

#### 2.3.3 Sparsification of the similarity matrices

Recent studies on manifold learning theories and spectral graph have shown that the scattered nearest neighbors of data points can effectively model local geometric structure [53, 63]. In graph regularized matrix factorization, the nearest neighbor graph can promote close lncRNAs (or miRNAs) to be sufficiently close to each other in the lncRNA space (or miRNA space) [46, 64]. That is, it can preserve the local geometries of the original data. In this study, the affinity graphs (*S*^*L*∗^ and *S*^*M*∗^) for lncRNAs and miRNAs are constructed using *p*-nearest neighbor graph, respectively. We set *p*=5, and the weight matrix *G*^*l*^ is generated based on the lncRNA similarity matrix *S*^*L*^ as follows:

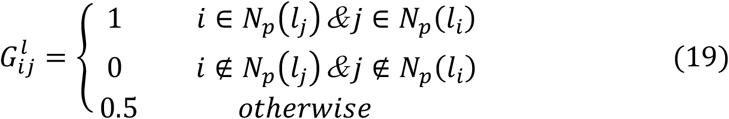

where *N*_*p*_(*l*_*i*_) and *N*_*p*_(*l*_*j*_) denote the sets of *p*-nearest neighbors to lncRNA *l*_*i*_ and lncRNA *l*_*j*_, respectively. Subsequently, the sparse similarity matrix *S*^*L*∗^ for lncRNAs is defined as:

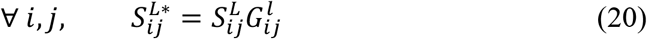

The same procedure for miRNAs, the sparse similarity matrix *S*^*M*∗^ can be obtained by the similarity matrix *S*^*M*^ of miRNA.

#### 2.3.4 The model of GNMFLMI

In the Euclidean space, the standard nonnegative matrix factorization in Eq. (2) fails to discover the intrinsic geometrical and discriminating structure of the data space [65, 66]. To avoid overfitting and enhance generalization capability, we use the Tikhonov (*L*_2_) regularization in Eq. (2) to guarantee the *U* and *V* smoothness (i.e. Eq. (6)) [55]. At the same time, graph regularization is used to ensure that the relative positions of data points in the lncRNA feature space or miRNA feature space are unchanged [46]. The objective function of graph regularized nonnegative matrix factorization can be defined as follows:

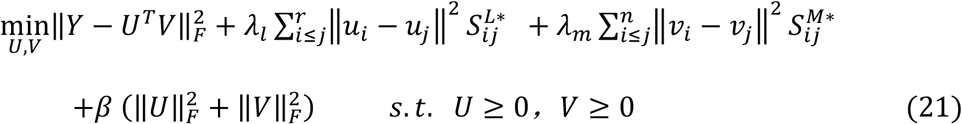

and

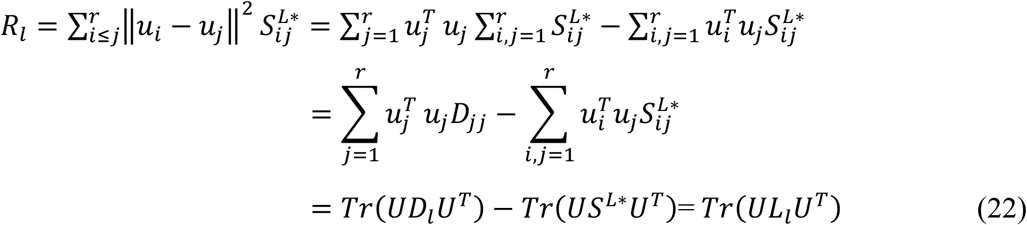

Similarly,

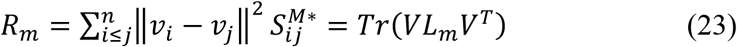

where *λ*_*l*_ and *λ*_*m*_ are the graph regularization parameters, *u*_*i*_ and *v*_*j*_ denote the *ith* and *jth* columns of *U* and *V*, respectively. *R*_*l*_ and *R*_*m*_ are the graph regularization terms of lncRNA and miRNA. We hope that if two data points *l*_*i*_ and *l*_*j*_ are close, *u*_*i*_ and *u*_*j*_ are also close to each other by minimizing *R*_*l*_ (or *R*_*m*_). *L*_*l*_ = *D*_*l*_ − *S*^*L*∗^ and *L*_*m*_ = *D*_*m*_ − *S*^*M*∗^ represent the graph Laplacian matrices for *S*^*L*∗^ and *S*^*M*∗^ [67], respectively; *D*_*l*_ and *D*_*m*_ are the diagonal matrices whose diagonal elements are column (or row) sums of *S*^*L*∗^ and *S*^*M*∗^, respectively. The Eq. (21) can be rewritten as:

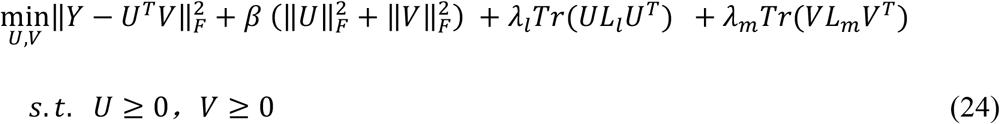

According to trace properties of matrix, the objective function Eq. (24) can be transformed into:

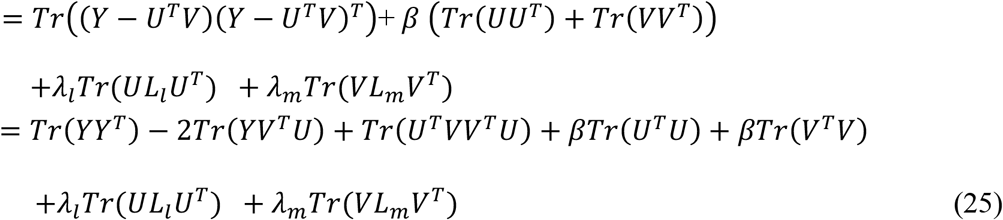

#### 2.3.5 Model optimization

Because of the objective functions of GNMFLMI is not convex, finding the global minima is unrealistic by optimization algorithm. However, the local minima can be achieved via algorithm. In this study, the Lagrange Multiplier method was introduced to solve the optimization problem in Eq. (25). Let *ψ* = {*φ*_*ki*_} and *Φ* = {*ϕ*_*kj*_}, the Lagrange multipliers *φ*_*ki*_ and *ϕ*_*kj*_ are used to restrict the *u*_*ki*_ ≥ 0 and *v*_*kj*_ ≥ 0, respectively. The Lagrange function ℋ can be constructed as:

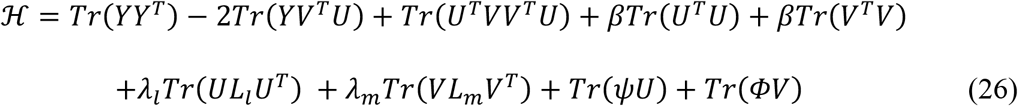

The partial derivatives of the function ℋ to *U* and *V* are:

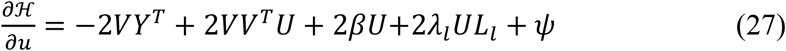

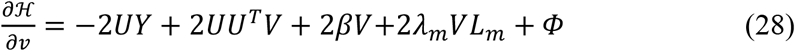

According to the Karush–Kuhn–Tucker (*KKT*) conditions [68], *φ*_*ki*_*u*_*ki*_ = 0 and *ϕ*_*kj*_*v*_*kj*_ = 0, we get the following equations for *u*_*ki*_ and *v*_*kj*_:

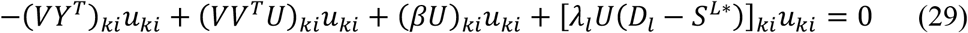

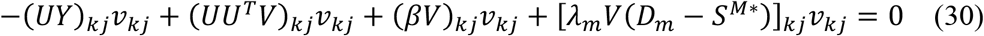

Finally, the updating rules can be determined as follows:

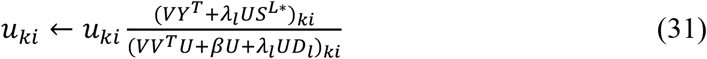

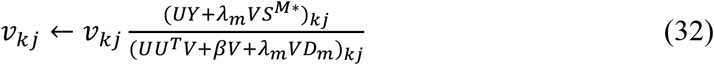

We update the nonnegative matrices *U* and *V* according to Eq. (31) and Eq. (32) until convergence or reaching the iteration upper limit. Ultimately, the new prediction lncRNA-miRNA interaction matrix *Y*^∗^ can be calculated by *Y*^∗^ = *U*^*T*^*V*. In general, the larger value of the element in prediction matrix *Y*^∗^, it is more likely that lncRNA/miRNA interacts with the corresponding miRNA/lncRNA. That is, for each miRNA, we can sort the lncRNA in descending order according to the value of the element, the top ranked lnRNAs in each column of *Y*^∗^ are more likely to be associated with the corresponding miRNA. The same is true for each lncRNA. Table 1 generalizes the procedure of GNMFLMI for predicting lncRNA-miRNA interactions.

**Table 1.**
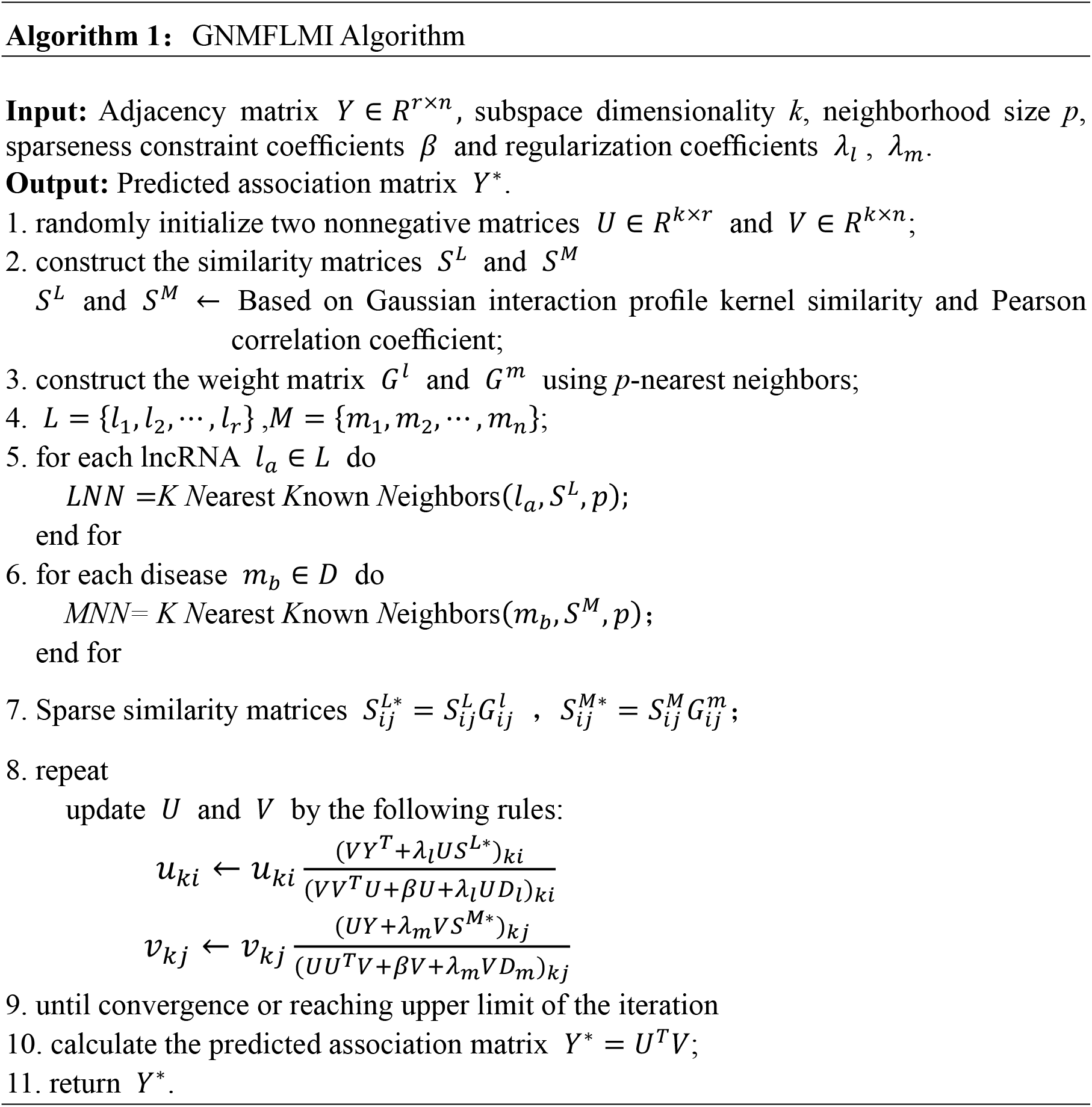
Algorithm description of graph regularized nonnegative matrix factorization for predicting lncRNA-miRNA interactions.

## 3 Results and discussion

### 3.1 Experimental settings

In this study, to estimate the performance of GNMFLMI on predicting lncRNA-miRNA interactions, the five-fold cross validation experiments were performed on the lncRNASNP2 dataset and compare our method with the following approaches: NMF and RNMF. In the five-fold cross validation, we randomly divide 8634 known lncRNA-miRNA interaction samples into five equal subsets. For each cross validation experiment, one of the subsets was used as the test set and the other four subsets were used as the training set.

The receiver operating characteristics (ROC) curve and AUC (area under the receiver operating characteristics curve) are widely used to estimate the performance [69, 70]. A larger value of AUC represents the better prediction performance of model. The confusion matrix can be obtained by setting different thresholds, the sensitivity (Sen.) and specificity (Spe.) are calculated as:

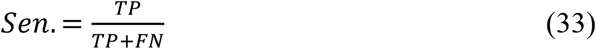

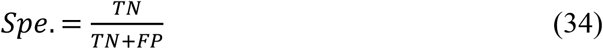

where FN and TP represent the number of false negative samples and true positive samples, respectively; FP and TN represent the number of false positive samples and true negative samples, respectively. FPR is false positive rate (FPR=1-Spe.), TPR is true positive rate (TPR=Sen.). In addition, precision (Pre.), accuracy (Acc.) and F1-Score are also used as general measurements.

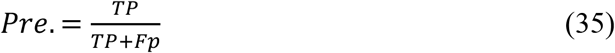

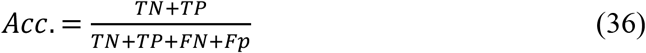

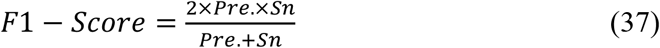

In this paper, the parameters are chose based on the grid search. There are five parameters in our method: neighborhood size *p*, subspace dimensionality *k*, sparseness constraint coefficient *β* and graph regularization coefficients *λ*_*l*_, *λ*_*m*_. The parameter combinations were determined from the following ranges: {65,140} for *k*, {0.0001,0.001,0.01} for *β* and the ranges of *λ*_*l*_ = *λ*_*m*_ ∈ {0.001,0.01,0.1}. Based on the studies of Cai *et al.* [46] and Li *et al.* [65], we set *p* = 5. Finally, the parameter values are *p* = 5, *k=*80, *β* = 0.01, *λ*_*l*_ = *λ*_*m*_ = 0.1.

### 3.2 Cross validation

We compared the performance of GNMFLMI with computational approaches NMF and CNMF on the lncRNASNP2 dataset. Figure 2, Figure 3 and Figure 4 plot ROC curves and calculate the average AUC values of NMF, CNMF and GNMFLMI, respectively. Table 2 lists the AUC values of GNMFLMI, CNMF and NMF under five-fold cross validation. GNMFLMI achieved the AUC value of 0.9769, which higher than the AUC values of NMF 0.9344 and CNMF 0.9510. The experiment results show that the prediction performance of GNMFLMI outperform the NMF and CNMF.

**Table 2.** The average AUC values and standard deviations obtained by various methods under five-fold cross validation (CV) on the lncRNASNP2 dataset.

| Methods | The AUC values under five-fold CV |  |  |  |  | Average |
| --- | --- | --- | --- | --- | --- | --- |
|  | 1st | 2nd | 3rd | 4th | 5th |  |
| NMF | 0.9399 | 0.9383 | 0.9267 | 0.9331 | 0.9341 | 0.9344±0.0052 |
| CNMF | 0.9562 | 0.9488 | 0.9476 | 0.9453 | 0.9572 | 0.9510±0.0054 |
| GNMFLMI | 0.9748 | 0.9752 | 0.9763 | 0.9799 | 0.9781 | 0.9769±0.0022 |

**Fig. 2.**
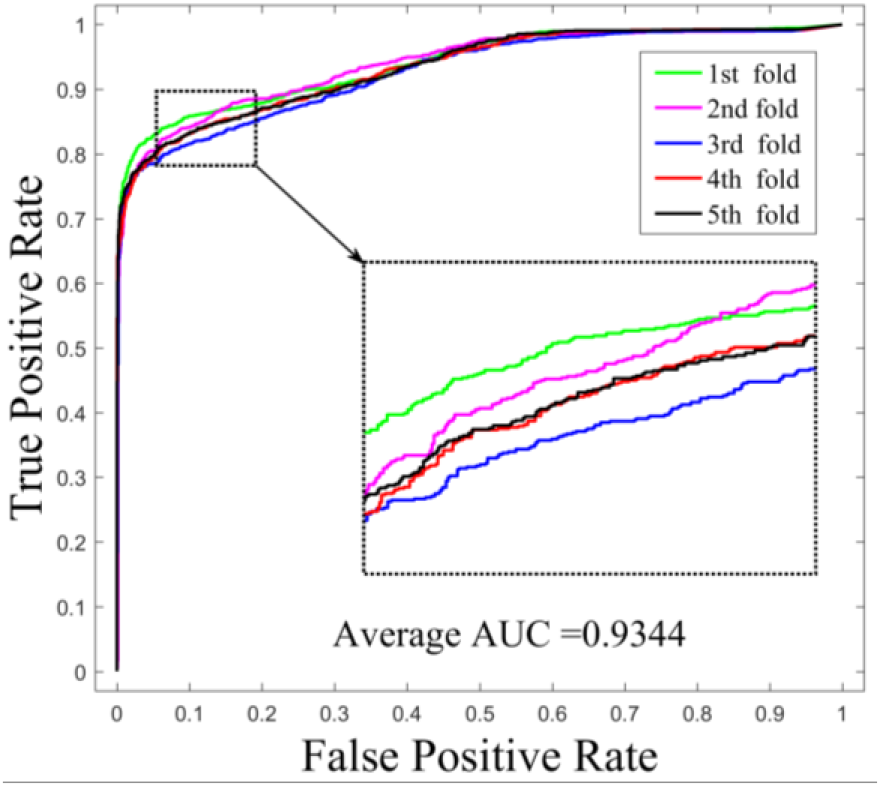
The ROC curves of NMF in lncRNA-miRNA interaction prediction under 5-fold cross validation.

**Fig. 3.**
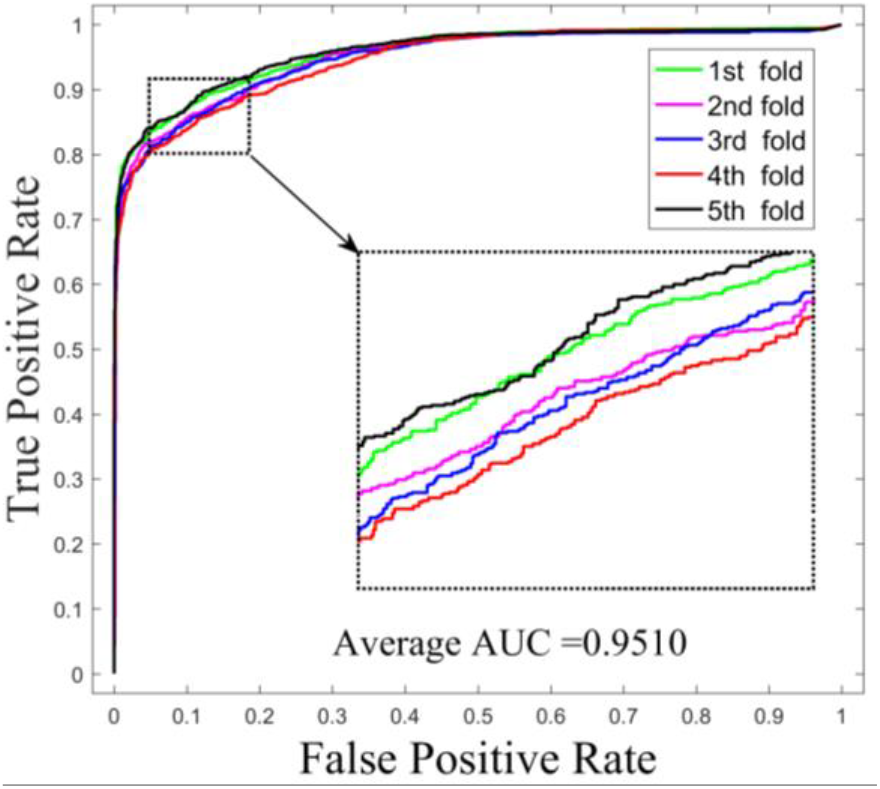
The ROC curves of CNMF in lncRNA-miRNA interaction prediction under 5-fold cross validation.

**Fig. 4.**
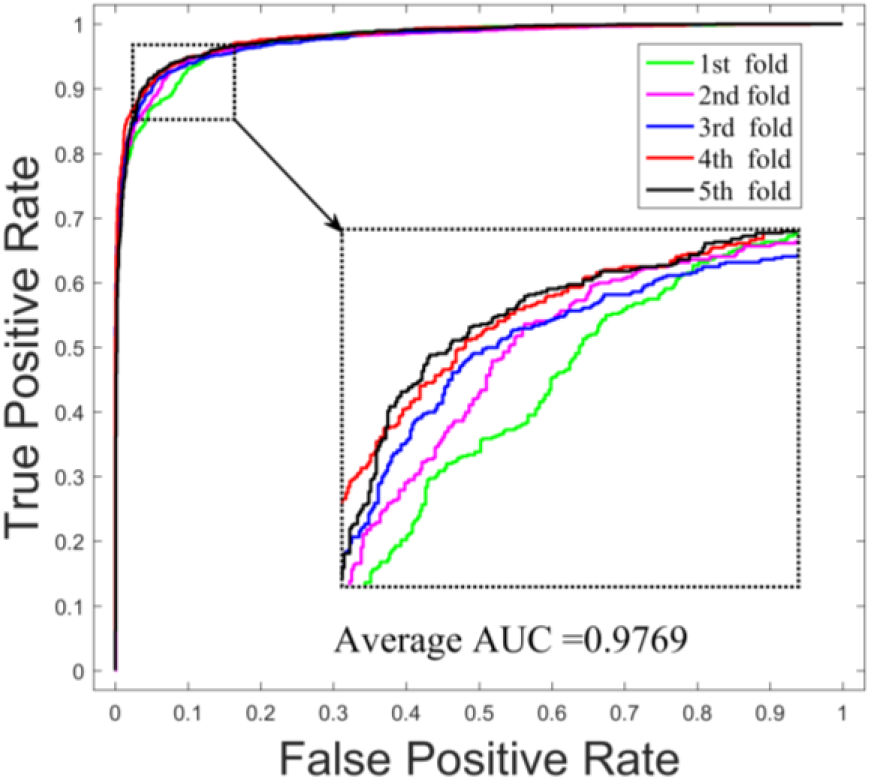
The ROC curves of GNMFLMI in lncRNA-miRNA interaction prediction under 5-fold cross validation.

In addition, the sensitivity, precision, accuracy and F1-Score for these methods were calculated at different specificity. As shown in Table 3, when specificity is 95%, the average sensitivities of GNMFLMI, NMF and CNMF are 89.40%, 80.17% and 82.08%, respectively. The sensitivity of GNMFLMI is 9.23% and 7.32% higher than NMF and CNMF. When specificity is 90%, GNMFLMI achieves the average sensitivity of 94.20%, which is still 10.63% and 8.47% higher than NMF and CNMF, respectively. In figure 5, we can also discover that the ROC curve of GNMFLMI is always above CNMF and NMF. These results further demonstrate that the performance of GNMFLMI is better than CNMF and NMF.

**Table 3.** The average sensitivity, precision, accuracy and F1-Score values of GNMFLMI and existing methods at different specificity.

| Methods | Spe.(%) | Sen.(%) | Pre.(%) | Acc.(%) | F1-Score(%) |
| --- | --- | --- | --- | --- | --- |
| NMF | 95.00 | 80.17 | 94.15 | 87.59 | 86.59 |
| CNMF | 95.00 | 82.08 | 94.28 | 89.46 | 88.83 |
| GNMFLMI | 95.00 | 89.40 | 94.72 | 92.21 | 91.98 |
| NMF | 90.00 | 83.57 | 89.29 | 86.78 | 86.33 |
| CNMF | 90.00 | 85.73 | 89.53 | 87.86 | 87.54 |
| GNMFLMI | 90.00 | 94.20 | 90.38 | 92.09 | 92.25 |

**Fig. 5.**
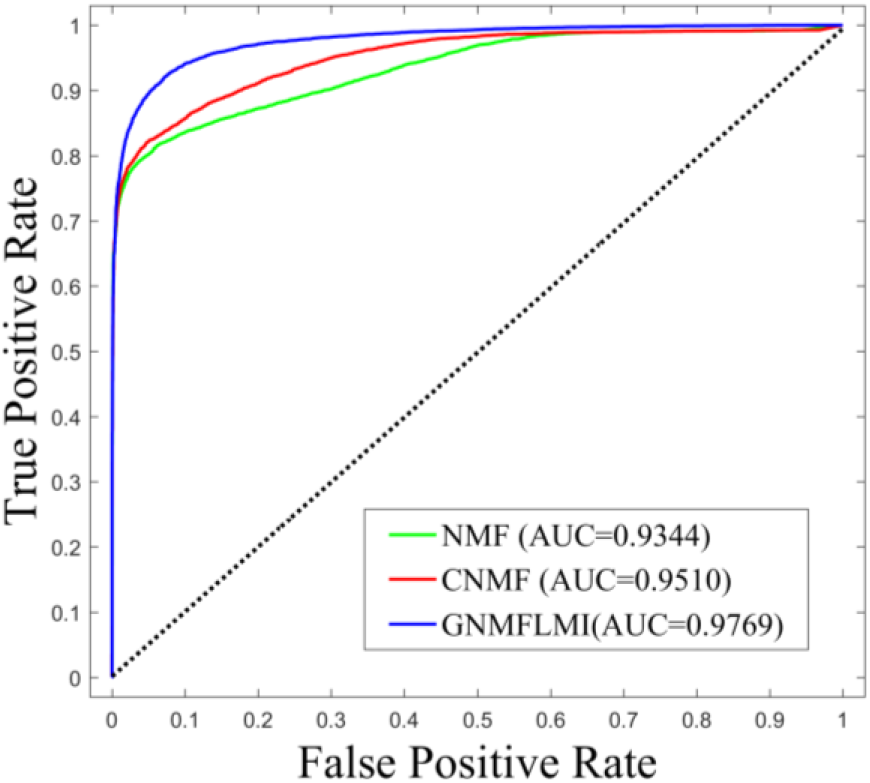
The ROC curves of three methods.

### 3.3 Case study

Case studies are carried out to further verify the capability of GNMFLMI on predicting novel interactions between lncRNAs and miRNAs. Here, we left out an arbitrary lncRNA-miRNA interaction (i.e. removing its interactions from the lncRNASNP2 dataset) to verify if its interactions would be discovered successfully. According to [71], the prediction performance is greatly affected by nearest neighbor information. If the new lncRNA *l*_*i*_ (miRNA *m*_*i*_) and lncRNA *l*_*j*_ (miRNA *m*_*j*_) are close to each other, it may be easy to accurately predict the interactions between them, and vice versa. In this work, we select the lncRNA and miRNA which the similarity to the nearest neighbor is low (according to *S*^*L*∗^ and *S*^*M*∗^, respectively) to validate the capability of model on predicting novel interactions. However, the standard NMF and CNMF fails to discover the novel interactions in this case.

For lncRNA-nonhsat159254.1, we remove all miRNAs which interact with this lncRNA from the lncRNASNP2 dataset, the remaining known interactions are used to train the model of GNMFLMI. Then, all candidate miRNAs are sorted in descending order according to the predicted interaction scores. Table 4 gives the top 20 predicted interactions for nonhsat159254.1, the top 20 predicted interactions were verified by databases. The same procedure is performed for miRNA-hsa-mir-544a, Table 5 lists the top 20 predicted interactions for hsa-mir-544a, 16 out of the top 20 candidate lncRNAs were verified by databases.

**Table 4.** The top 20 novel interactions predicted by GNMFLMI for nonhsat-159254.1 on the lncRNASNP2 dataset.

| Rank | miRNAs | Evidences |
| --- | --- | --- |
| 1 | hsa-mir-590-3p | confirmed |
| 2 | hsa-mir-150-5p | confirmed |
| 3 | hsa-mir-374b-5p | confirmed |
| 4 | hsa-mir-374a-5p | confirmed |
| 5 | hsa-mir-30d-5p | confirmed |
| 6 | hsa-mir-30e-5p | confirmed |
| 7 | hsa-mir-30a-5p | confirmed |
| 8 | hsa-mir-30c-5p | confirmed |
| 9 | hsa-mir-30b-5p | confirmed |
| 10 | hsa-mir-200a-3p | confirmed |
| 11 | hsa-mir-141-3p | confirmed |
| 12 | hsa-mir-205-5p | confirmed |
| 13 | hsa-mir-216a-5p | confirmed |
| 14 | hsa-mir-4465 | confirmed |
| 15 | hsa-mir-26a-5p | confirmed |
| 16 | hsa-mir-1297 | confirmed |
| 17 | hsa-mir-26b-5p | confirmed |
| 18 | hsa-mir-363-3p | confirmed |
| 19 | hsa-mir-25-3p | confirmed |
| 20 | hsa-mir-92a-3p | confirmed |

**Table 5.** The top 20 novel interactions predicted by GNMFLMI for hsa-mir-544a on the lncRNASNP2 dataset.

| Rank | LncRNAs | Evidences |
| --- | --- | --- |
| 1 | nonhsat137542.2 | confirmed |
| 2 | nonhsat137541.2 | confirmed |
| 3 | nonhsat137559.2 | confirmed |
| 4 | nonhsat159243.1 | confirmed |
| 5 | nonhsat137558.2 | confirmed |
| 6 | nonhsat022125.2 | confirmed |
| 7 | nonhsat159246.1 | confirmed |
| 8 | nonhsat159242.1 | confirmed |
| 9 | nonhsat159254.1 | confirmed |
| 10 | **nonhsat096369.2** | **unconfirmed** |
| 11 | **nonhsat096376.2** | **unconfirmed** |
| 12 | **nonhsat096375.2** | **unconfirmed** |
| 13 | nonhsat022145.2 | confirmed |
| 14 | nonhsat159244.1 | confirmed |
| 15 | nonhsat159252.1 | confirmed |
| 16 | **nonhsat198591.1** | **unconfirmed** |
| 17 | nonhsat159248.1 | confirmed |
| 18 | nonhsat022132.2 | confirmed |
| 19 | nonhsat007675.2 | confirmed |
| 20 | nonhsat007699.2 | confirmed |

It is worth noting that the nonhsat159254.1 and hsa-mir-544a prediction can be considered two difficult cases. Specifically, the similarities both nonhsat159254.1 and hsa-mir-544a to their nearest neighbors lncRNA and miRNA are as low as 0.1786 (according to *S*^*L*∗^) and 0.3658 (according to *S*^*M*∗^), respectively. Such low similarity makes it more difficult to predict the interactions between them. According to the above two cases, it is shown that GNMFLMI can effectively predict novel and challenging interactions between lncRNAs and miRNAs, which can provide valuable information for biological experiments

## 4 Conclusion

The interactions between lncRNAs and miRNAs constitute a complex molecular regulatory network, and studies have confirmed that their interactions are closely related to the occurrence and development of various diseases. Identifying lncRNA-miRNA interactions can help people better understand the complex disease mechanisms. In this paper, we propose a new method, GNMFLMI, for lncRNA-miRNA interaction prediction. Different from other traditional methods, GNMFLMI guides the matrix factorization via constructing graph Laplacian regularizations of lncRNAs and miRNAs, and uses Lagrange multipliers method to optimize the objective function. This method can also be applied into other similar association prediction (e.g. small molecular-miRNA and mRNA–protein associations). Five-fold cross validation and case studies were conducted to validate the performance of GNMFLMI, the experiment results show that our method outperforms the other compared methods and can identify potential lncRNA-miRNA interactions. We believe that our approach is helpful for clinical research. In future work, more different biological information can be integrated to further improve prediction performance of model.

## Acknowledgements

This work was supported in part by the NSFC Excellent Young Scholars Program, under Grant 61722212, in part by the National Natural Science Foundation of China, under Grants 61702444, 61572506, in part by the Pioneer Hundred Talents Program of Chinese Academy of Sciences, in part by the Chinese Postdoctoral Science Foundation, under Grant 2019M653804, in part by the West Light Foundation of the Chinese Academy of Sciences, under Grant 2018-XBQNXZ-B-008.

## Competing interests

The authors declare that they have no competing interests.

